# Analysis of P granules *in vivo* and *ex vivo*

**DOI:** 10.1101/245878

**Authors:** Andrea Putnam, Madeline Cassani, Jarrett Smith, Geraldine Seydoux

## Abstract

*Corrected version -This version can be cited*. RNA granules are dynamic sub-cellular compartments that lack enveloping membranes. RNA granules have been proposed to form by liquid-liquid phase separation, a thermodynamic process that partitions molecules between dilute and condensed liquid phases ^1^. P granules are archetypal RNA granules in *C. elegans* that display liquid-like behaviors ^2^. Here we describe *in vivo* and *ex vivo* approaches to analyze the material properties of P granules. We find that the liquid phase of P granules is stabilized by a molecularly-distinct, enveloping shell that is intrinsically non-dynamic. Consistent with a gel phase, the shell is resistant to dilution, high salt, and aliphatic alcohols, and dissolves in SDS. Solidification of RNA granules has been linked to neuronal degeneration ^3^. Our findings suggest that gel-like polymers are essential components of RNA granules that help stabilize liquid phases in the cellular environment.

## MAIN TEXT

Liquid-liquid phase separation (LLPS) is a spontaneous de-mixing process where molecules partition between condensed and dilute liquid phases ^1,4^. LLPS has been proposed to underlie the formation of RNA granules in cells, and several RNA-binding proteins have been found to undergo LLPS *in vitro* ^5^. Some LLPS condensates assembled *in vitro* mature over time into non-dynamic, gel-like condensates ^6-9^. Solidification of liquid phases in RNA granules has been linked to pathological states, as in certain neurodegenerative diseases where RNA granule components become trapped in non-dynamic aggregates ^4,10^. Solid cores have also been observed in stress granules, but whether they are by-products of LLPS or essential structures that support granule assembly is not yet known ^11,12^. Here, we have investigated a possible role for non-dynamic condensates in the formation of P granules, RNA granules that form under normal conditions in the germline of *C. elegans*.

P granules were the first cytoplasmic RNA granules reported to display liquid-like behaviors ^2^. P granules are roughly round, can fuse with each other, and exchange components with the cytoplasm. P granules are perinuclear during most of germline development and become cytoplasmic at the oocyte-to-embryo transition ^13^. During the first embryonic cell cycle, P granules disassemble and reassemble in the germ plasm, a region of cytoplasm in the posterior of the embryo that is partitioned to the germ lineage ^2^ (Fig. 1). Genetic analyses have identified two classes of proteins required for P granule assembly in embryos: the RGG-domain protein PGL-3 (and its paralog PGL-1), which form condensates that recruit other RNA-binding proteins ^14,15^, and the intrinsically-disordered protein MEG-3 (and its paralog MEG-4), which localizes and stabilizes PGL condensates in germ plasm ^16,17^. PGL-3 and MEG-3 overlap only partially in P granules, with PGL-3 enriched in the center and MEG-3 enriched at the periphery ^17^. Lattice light sheet microscopy revealed that MEG-3 forms a structured shell around a more amorphous PGL core, raising the possibility that the MEG and PGL phases have distinct material properties ^17^.

**Figure 1.**
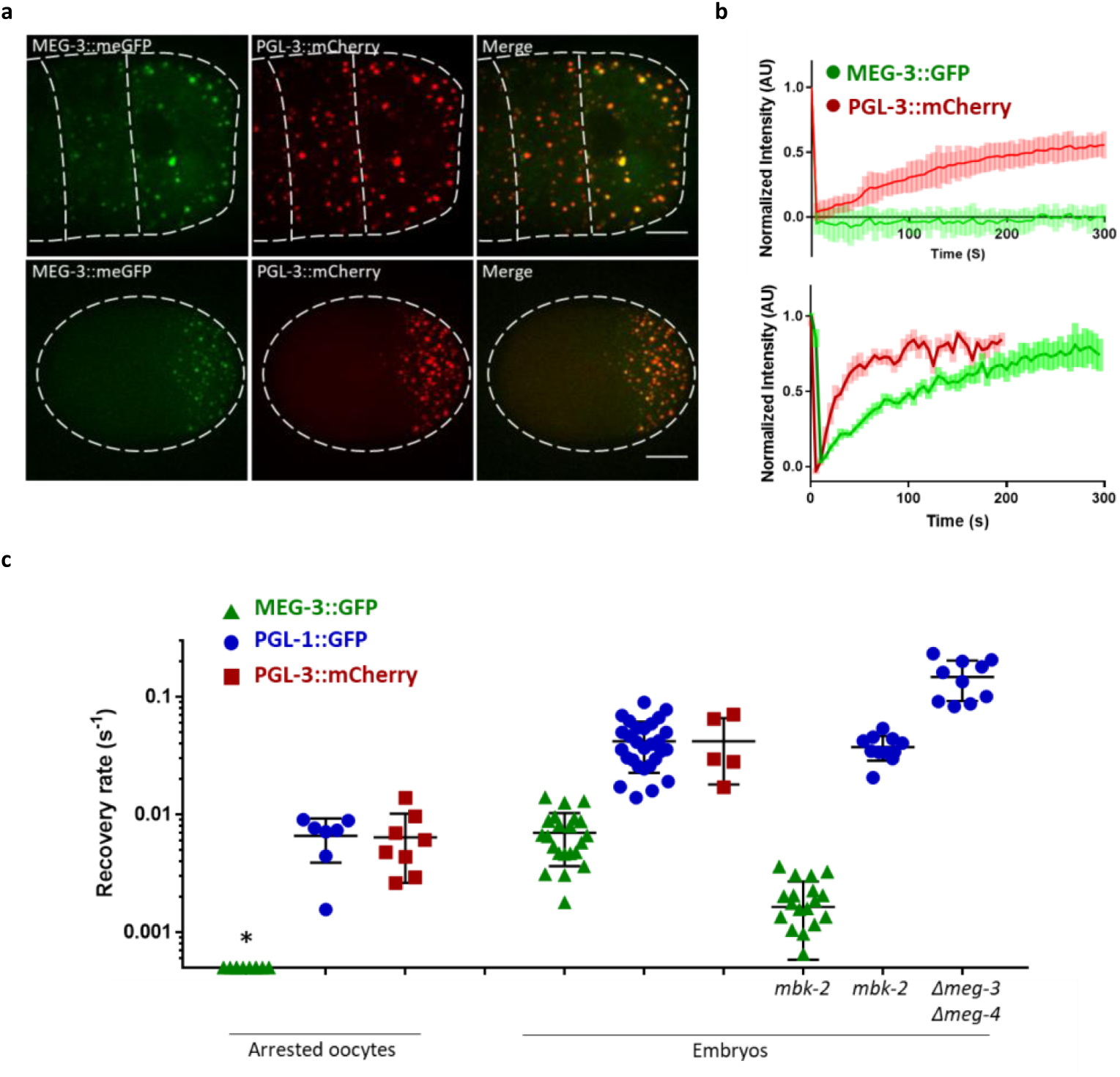
P granules contain two phases that exhibit distinct dynamics. (A) Photomicrographs of arrested oocytes (top row) and a 1-cell embryo (bottom row) expressing MEG-3::meGFP and PGL-3::mCherry fusions (strain JH3602). In this and all other figures, embryos are oriented with its posterior end (germ plasm) to the right. (B) Graphs showing MEG-3::meGFP or PGL-3::mCherry fluorescence recovery after photobleaching (FRAP) in arrested oocytes or embryos. Granule intensity was measured every 5s for 300s before and after bleaching. Values were normalized to initial fluorescence intensity and plotted as an average from multiple oocytes or embryos. Error bars represent mean ± SEM. (C) Graphs showing recovery rates from FRAP experiments plotted in B or SF1B. Each symbol represents one FRAP experiment on one granule. The rate of fluorescence recovery (Methods) is plotted on the Y axis on a log scale. The X axis indicates the stage and genotype tested. MEG-3 dynamics are slower than PGL-1 and PGL-3 dynamics under all conditions. Error bars represent mean ± SD. For data directly on the x-axis of each graph indicated by a “*”, no recovery was measured.

To characterize the properties of the MEG and PGL phases, we tagged the *meg-3*, *pgl-1* and *pgl-3* loci with GFP or mCherry using CRISPR genome editing (Fig. 1a and ^16,18^). We used fluorescence recovery after photobleaching (FRAP) to determine rates of cytoplasm to granule exchange at two developmental stages: in arrested oocytes where P granules are stable, and in embryos where P granules undergo cycles of dissolution and condensation as they segregate with the germ plasm (Supplemental Movie 1). In arrested oocytes, MEG-3 forms stable condensates that were non-dynamic over the time scale tested (5 minutes), whereas PGL-1 and PGL-3 exchanged at a rate of 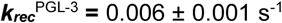 (Fig. 1b,c). In embryos, MEG-3 became dynamic, but remained significantly slower than PGL-1 and PGL-3 whose dynamics also increased 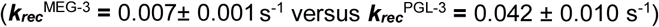 (Fig. 1b,c). MBK-2 is a kinase that becomes activated during the oocyte-to-embryo transition to promote P granule dissolution ^17,19^. We found that MBK-2 is required to stimulate MEG-3 dynamics in embryos: MEG-3 dynamics were significantly slower in embryos depleted of MBK −2 by RNAi 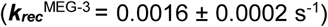 (Fig. 1c, Extended Data Fig. 1b). PGL-1 dynamics, in contrast, were not affected by loss of MBK-2, but were affected by the loss of *meg-3* and *meg-4* (Fig. 1c, Extended Data Fig. S1b). PGL dynamics were significantly faster in *meg-3meg-4* mutants 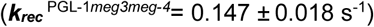 compared to wild-type 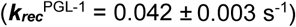 (Fig. 1C, Extended Data Fig. 1b). In *meg-3meg-4* mutants, PGL condensates are fewer, do not localize to the posterior and do not persist past the 4-cell stage (Extended Data Fig. 1a and Supplemental Movie 2). These observations confirm that P granules contain two phases with distinct properties: a MEG-3 phase that becomes dynamic in embryos, and a constitutively dynamic, possibly liquid, PGL phase that is stabilized by the MEG-3 phase.

Liquid phases can be sensitive to changes in temperature and concentration ^20,21^. To test the effect of temperature, we filmed embryos expressing PGL-3::mCherry and MEG-3::meGFP while subjecting them to rapid temperature shifts using a temperature-controlled stage. An up-shift from 20°C (normal growth temperature) to 34°C caused PGL-3::mCherry to dissolve into the cytoplasm within seconds. Return to 20°C caused PGL-3::mCherry to reappear on the granules after ~ 8 min (Fig. 2a). MEG-3::meGFP, in contrast, remained in granules throughout the experiment. A longer 15 minute shift to 34°C also did not affect MEG-3::meGFP distribution (Extended Data Fig. 2a). To test the effect of concentration, we punctured the eggshell of the embryo with a laser to release cellular contents into an aqueous buffer (Fig. 2b – See Methods for a detailed protocol that avoids technical artifacts that confounded initial experiments reported in the previous versions of this manuscript). Consistent with a liquid phase, PGL-3::mCherry diffused immediately upon dilution into the buffer. PGL-1::GFP also dissolved immediately *ex vivo* (Extended Data Fig. 2b). MEG-3::meGFP, in contrast, persisted in the granule phase, as the granules flowed out of the embryo into the dilute buffer (Fig. 2b and Supplemental Movie 3). On average, 64 ± 18% of MEG-3::meGFP fluorescence remained in the granule phase after extrusion (Fig. 2c). The extruded MEG-3::meGFP granules persisted in buffer for at least 1 hour (Extended Data Fig. 2c,d). MEG-3::meGFP granules remained stable when extruded into buffer containing 1M NaCl, 1,6 hexanediol (5%), or 10mM ATP, chemicals that dissolve liquid phases ^22,23^ (Fig. 2c). MEG-3::meGFP granules, however, dissolved in SDS (0.5%) (Fig. 2c). These observations suggest that the MEG-3 phase is not an irreversible aggregate, but a stable, gel-like phase. In contrast, the PGL phase behaves like a liquid.

**Figure 2.**
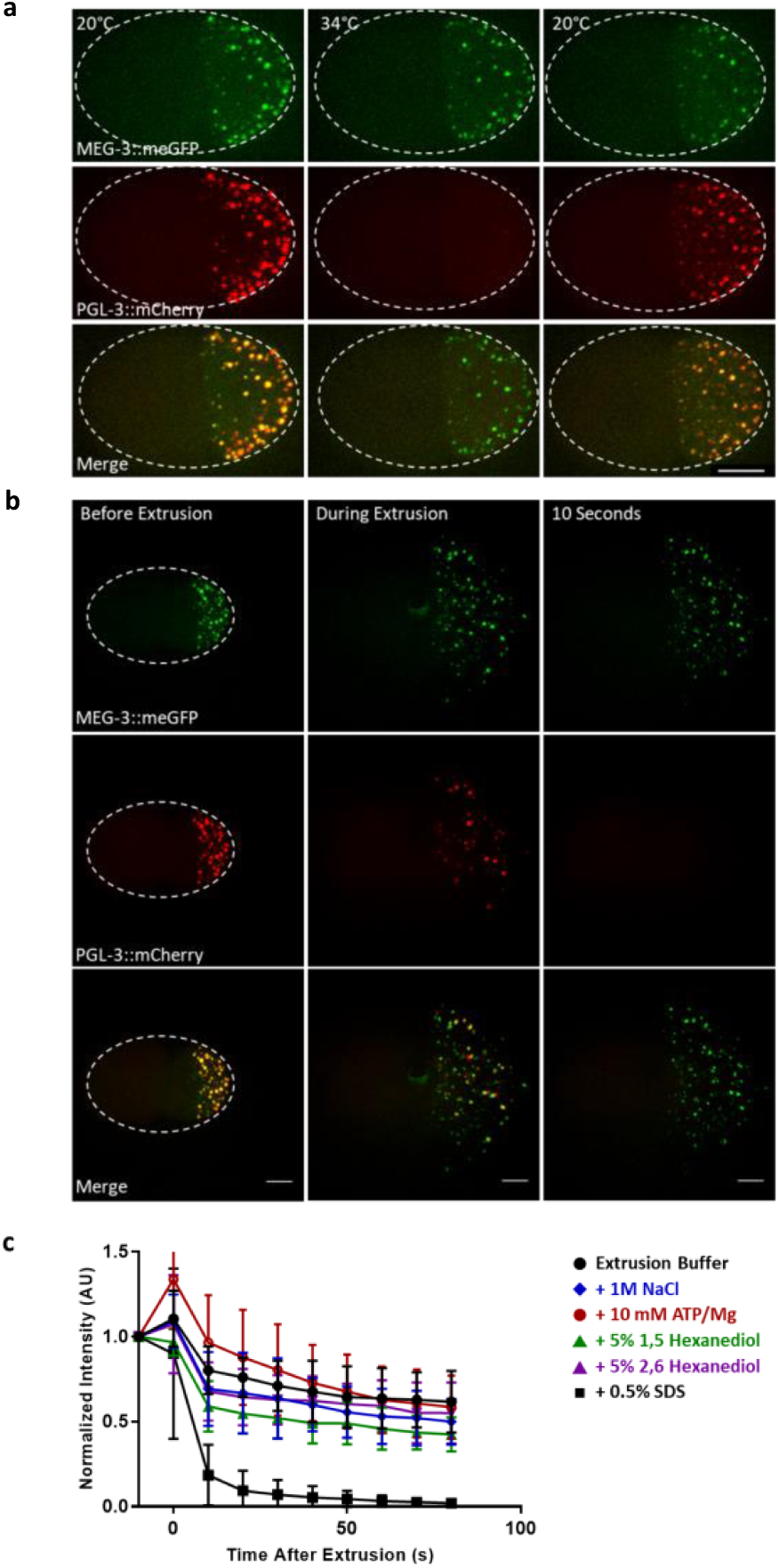
Effect of temperature and dilution on the MEG-3 and PGL-3 phases. (A) Time-lapse photomicrographs of a two-cell embryo expressing MEG-3::meGFP and PGL-3::mCherry fusions (JH3602) and cultured at 20°C, up-shifted to 34°C for 1 minute, and downshifted back to 20°C for 8 minutes. MEG-3 remains in granules throughout. PGL-3 disperses at 34°C and returns to P granules at 20°C. Scale bar is 10 μm. (B) Time-lapse photomicrographs of a two-cell embryo expressing MEG-3::meGFP and PGL-3::mCherry fusions (JH3602) before, immediately after (~1s), and 10s after laser puncture of the eggshell. Laser puncture causes the contents of the posterior blastomere to spill onto the slide mixing with the aqueous buffer. MEG-3::GFP persists in the granule phase, whereas mCherry::PGL-3 becomes dispersed. Scale bar is 10 μm. (C) Graph showing the fraction of MEG-3::GFP retained in the granular phase after extrusion. Total GFP fluorescence from granules was measured before laser puncture (IB) and after laser puncture (IA). Means are indicated along with error bars representing ± SD.

P granules were the first RNA granules proposed to behave like liquid droplets suspended in the cytoplasm. Our observations indicate that the liquid phase of P granules is stabilized by a molecularly distinct enveloping layer that has gel-like properties. Consistent with a gel-like phase, the MEG-3 shell is non-dynamic, resistant to dilution, high salt and aliphatic alcohol, and sensitive to SDS. MEG-3 is essential to assemble PGL droplets in the germ plasm ^17^. The MEG-3 shell may therefore function as a protective “skin” that stabilizes the liquid core of P granules against factors in the cytoplasm that would otherwise dissolve the PGL phase. In embryos where P granules initiate cycles of disassembly and reassembly, the MEG-3 phase becomes dynamic in a manner that depends on DYRK/MBK-2 kinase. Assemblies that transition between gel and liquid-like dynamics have also been suggested to exist in nucleoli and stress granules, which solidify *ex vivo and* exhibit ATP-dependent dynamics *in vivo* ^11,24^. Solidification of RNA granules has been linked to aging, ATP-deprived environments and certain neurodegenerative disorders ^4,10^. Our findings suggest that gel phases are natural and essential stabilizing components of RNA granules that, like polymers in other self-organizing structures (e.g. cytoskeleton), can cycle between dynamic and non-dynamic states ^25,26^. Models that only consider equilibrium dynamics therefore, may not be sufficient to describe the properties of RNA granules in cells.

## MATERIALS AND METHODS

### Strain maintenance and RNA-mediated interference

*C. elegans* was cultured according to standard methods at 20°C unless otherwise stated ^27^. Genome editing was performed using CRISPR/Cas9 as previously described ^28^. Strains used in this study are listed in Table S1.

*mbk-2(RNAi)* was performed by feeding ^29^ using plasmids from the Ahringer or Openbiosystems libraries ^17^. HT115 bacteria transformed with feeding vectors were grown at 37°C in LB + ampicillin (100 µg/ml) for 5 hr, induced with 5 mM IPTG for 45 min, plated on NNGM (nematode nutritional growth media) + ampicillin (100 µg/ml) + IPTG (1 mM), and grown overnight at room temperature. Embryos were hatched on RNAi plates and allowed to feed for ~3-4 days

For visualization of arrested oocytes, *fog-2(q71)* females, which do not produce sperm, were collected in the L4 stage and matured for 8 hours before imaging in 20 nM levamisole. For imaging of embryos, adult females were dissected in 1X *ex vivo* buffer (25 mM HEPES pH 7, 150 mM NaCl, 2 mM CaCl_2_, 2 mM MgCl_2_) to release embryos which were mounted onto 2% agarose pads on glass slides.

### Microscopy

Images were captured with a Zeiss Axio Imager equipped with a Yokogawa spinning-disc confocal scanner and Slidebook v 6.0 software (Intelligent Imaging Innovations). Unless otherwise noted embryo and oocyte images are Z stack maximum projections using a Z step size of 1 μm, spanning the depth of the oocyte or embryos using a 60x objective. All image analyses were conducted using the Fiji image-processing package (http://fiji.sc/Fiji).

### Fluorescence Recovery after Photobleaching (FRAP)

Photobleaching of MEG-3::meGFP and PGL-3::mCherry in oocytes or embryos was performed using a laser to bleach a region slightly larger than the area of the granule (2 μm diameter) in the center plane of the Z-stack. Images were acquired as Z-stacks with 10 planes with a step size 1 µm (oocytes) or 11 planes with a step size of 0.6 µm (embryos) by imaging 50ms exposures in the 488 channel and 25ms (embryos) or 50ms (arrested oocytes) exposures in the 561 channel using a 63x objective. Images were acquired every 3 or 5s during a recovery phase of 180-300s after photobleaching.

Maximum projections of Z-stacks acquired as above were analyzed in Fiji. The total or mean fluorescence intensity of the area containing the granule was measured at each time point. Fluorescence recovery was corrected for background and normalized to the initial granule intensity using the equation: n*I* = (*I-I^bkg^*)/(*I^i^*-*I^bkgi^*), where n*I* is the background corrected and normalized fluorescence intensity, *I* is the intensity of the FRAPed granule, *I^bkg^* is the fluorescence intensity of the unFRAPed cytoplasm, *I^i^* is the initial intensity of the FRAPed granule before bleaching, and *I^bkgi^* is the initial background intensity. For MEG-3::meGFP FRAP experiments in *mbk-2(RNAi)* embryos where amplitudes only just approached full recovery during the time frame of the experiment, final recovery amplitudes were limited to A^rec^ = 1.

### Temperature shifts

Temperature shift experiments were performed on a fluidic temperature control stage which enabled rapid temperature shifts (CherryTemp™, CherryBiotech). Embryos were prepared as described for FRAP experiments. Images were acquired using a z step size of 1µm, spanning the entire depth of the embryo. Final images are Z-stack maximum projections. Images were taken every 15s broken into the following segments: 1 min at 20°C followed by a 3-15 min shift to 34°C and then returned to 20°C for 25 min.

### *Ex vivo* Experiments

All *ex vivo* buffers were prepared in 25 mM HEPES pH 7, 150 mM NaCl, 2 mM CaCl_2_, 2 mM MgCl_2_ buffered to pH 7.0 with NaOH and HCl using a Mettler Toledo Seveneasy pH meter and a Fisherbrand™ Accumet™ glass body standard combination mercury free electrode. Embryos were dissected from adult hermaphrodites in *ex vivo* buffer and mounted on glass slides with 3% agarose pads made with 1X *ex vivo* buffer. Embryo contents were extruded by puncturing the eggshell near the anterior region of the germline blastomere using a 3i Ablate!™ laser system at 532 nm pulse setting with a power level of 155.

Note: to generate reproducible results, we have found that it is essential to mount embryos on 3% agarose pads made with 1X *ex vivo* buffer, and to puncture the embryo at the edge of the anterior region of the germline blastomere. Aiming the laser too close to the P granules gave inconsistent results, possibly due to the heat generated from the laser. We also confirmed this by manually extruding P granules on a glass slide, without the use of the laser (data not shown). It is also important to verify the pH of the buffer after addition of any additives, such as nucleotides.

All embryo images are Z stack maximum projections using a Z step size of 1 μm, spanning the entire depth of the embryo. Images were acquired using 50ms exposures in the 488 channel and/or 50ms exposures in the 561 channel every 10s using a 63x objective. For Extended Data Fig.2b, images were taken every 15min for 60min following extrusion.

To quantify MEG-3::meGFP persistence in granules, photomicrographs acquired as described above were analyzed using ImageJ64. Background was subtracted from each image using a rolling ball radius of 50 pixels. Pixels were smoothed 1 time and a pixel brightness threshold was set to 50-255 and the total integrated density of the objects (MEG-3::meGFP granules) was quantified. Total fluorescence intensity was calculated before (IB) and after (IA) extrusion and used to calculate a fluorescence ratio (IA/IB). For some embryos, granules left the field of view and could not be counted. The IA /IB, therefore, is a minimal estimate of the fraction of MEG-3::meGFP that remains in the granule phase after extrusion.

### Graphing and data fitting

All data was plotted and statistical analysis was conducted using Graphpad Prism 7 software. Fitting of recovery curves in FRAP experiments was conducted using Kaleidagraph (Synergy) software.

## ACKNOWLEDGEMENTS

We thank Dominique Rasoloson who provided strains JH3269 and JH3379 and Helen Schmidt who provided strain JH3606, and Andrew Folkmann for comments on the manuscript. Some strains were provided by the CGC, which is funded by NIH Office of Research Infrastructure Programs (P40 OD010440). This work was supported by the National Institutes of Health (NIH) (grant number R37 HD37047). G.S. is an investigator of the Howard Hughes Medical Institute.

## AUTHOR CONTRIBUTIONS

A.P, M.C, J.S, and G.S designed the research. A.P and M.C. performed all experiments, collected, and analyzed data. G.S and A.P prepared the manuscript with contributions from all authors.

## AUTHOR INFORMATION

The authors declare no competing interests. Correspondence and requests for materials should be addressed to G.S.

